# Liver-Chip: Reproducing Human and Cross-Species Toxicities

**DOI:** 10.1101/631002

**Authors:** Kyung-Jin Jang, Monicah A. Otieno, Janey Ronxhi, Heng-Keang Lim, Lorna Ewart, Konstantia Kodella, Debora Petropolis, Gauri Kulkarni, Jonathan E. Rubins, David Conegliano, Janna Nawroth, Damir Simic, Wing Lam, Monica Singer, Erio Barale, Bhanu Singh, Manisha Sonee, Anthony J. Streeter, Carl Manthey, Barry Jones, Abhishek Srivastava, Linda C. Andersson, Dominic Williams, Hyoungshin Park, Riccardo Barrile, Josiah Sliz, Anna Herland, Suzzette Haney, Katia Karalis, Donald E. Ingber, Geraldine A. Hamilton

## Abstract

Nonclinical rodent and non-rodent toxicity models used to support clinical trials of candidate drugs may produce discordant results or fail to predict complications in humans contributing to drug failures in the clinic. Here we applied microengineered Organ-on-Chip (Organ-Chip) technology to design rat, dog, and human Liver-Chips containing species-specific primary hepatocytes interfaced with liver sinusoidal endothelial cells, with or without Kupffer cells and hepatic stellate cells, cultured under physiological fluid flow. The Liver-Chips detected diverse phenotypes of liver toxicity including hepatocellular injury, steatosis, cholestasis, and fibrosis as well as species-specific toxicities when treated with tool compounds. Multi-species Liver-Chips may provide a useful platform for prediction of liver toxicity and inform human relevance of liver toxicities detected in animal studies to better determine safety and human risk.

**One Sentence Summary:** Microengineered Organ-Chip technology has been used to design rat, dog and human Liver-Chips that recapitulate species-specific liver toxicities.

---

The U.S. Food and Drug Administration (FDA) and European Medicines Agency generally require the safety of new drug candidates to be evaluated in both a rodent and a non-rodent animal models, frequently rat and dog, before moving the new chemical entity (NCE) into human clinical trials. An analysis of 150 drugs that caused adverse events in humans found that regulatory testing in rats and dogs correctly predicted just 71% of toxicities in humans (1). Moreover, while gastrointestinal, hematological, and cardiovascular toxicities were predicted with a relatively high concordance, the ability to predict liver toxicities was much lower. This was further confirmed by a more recent survey comparing target organ toxicities in animal and first-in-human studies that also found a low concordance of liver toxicity between human and animals (2). The poor prediction of liver toxicity in humans is driven by poor nonclinical to clinical translation and by rare idiosyncratic events that occur in large patient trials or at post-marketing. Thus, one of the major challenges the pharmaceutical and biotechnology industries face is selecting compounds with reduced risk for hepatotoxicity—the major cause for liver failure and drug attrition (3–6). Given the scale of this challenge and its negative impact on healthcare costs and development of new therapeutics, there is a critical need for more predictive and human relevant alternatives to animal models. Here, we explored whether human microengineered Organ-on-Chip (Organ-Chip) technology, which has been shown to faithfully recapitulate the complex functions and pathophysiology of multiple human organs (7–11), may be used to design species-specific Liver-Chips that can be used to address this challenge.

## Development and Characterization of rat, dog, and human Liver-Chips

Species-specific Liver-Chips lined by living rat, dog, or human hepatic cells were constructed using Organ-Chips that have previously been shown to recapitulate the multicellular architecture, tissue-tissue interface, vascular perfusion, fluid flow, and relevant physical microenvironment of multiple human organs, including lung, intestine, and kidney (10–12). Primary rat, dog, or human hepatocytes were seeded in the upper parenchymal channel within an extracellular matrix (ECM) sandwich (13) on top of an ECM-coated, porous membrane that separates the two parallel microchannels. Relevant species-specific rat, dog, or human liver sinusoidal endothelial cells (LSECs), with or without liver Kupffer cells and/or stellate cells, were cultured on the opposite side of the same membrane in the lower vascular channel (Fig. 1A). We initiated these studies by analyzing dual-cell Liver-Chips containing only the hepatocytes and LSECs (fig. S1A), which revealed that all three species of primary hepatocytes formed characteristic branched bile canalicular networks lined by functional multidrug resistance-associated protein 2 (MRP2) efflux transporters and maintained their stereotypical *in vivo*-like liver epithelial morphologies and cytoarchitecture for at least 14 days in culture under continuous flow (fig. S1B). In contrast, the same human, dog, and rat hepatocytes failed to form well developed bile canaliculi when maintained for 14 days without endothelium in the conventional static ECM sandwich culture plates (fig. S1B). It has been reported that sandwich culture can also form extensive canalicular network, but these are not sustained over 14 days and generally only over 7 days in culture (14, 15). In the microengineered Liver-Chip, the underlying vascular channel containing LSECs also displayed the multifunctional scavenger receptor stabilin-1, which is expressed selectively on sinusoidal endothelial cells of liver, spleen, and lymph nodes (fig. S1B) (16).

**Fig. 1.**
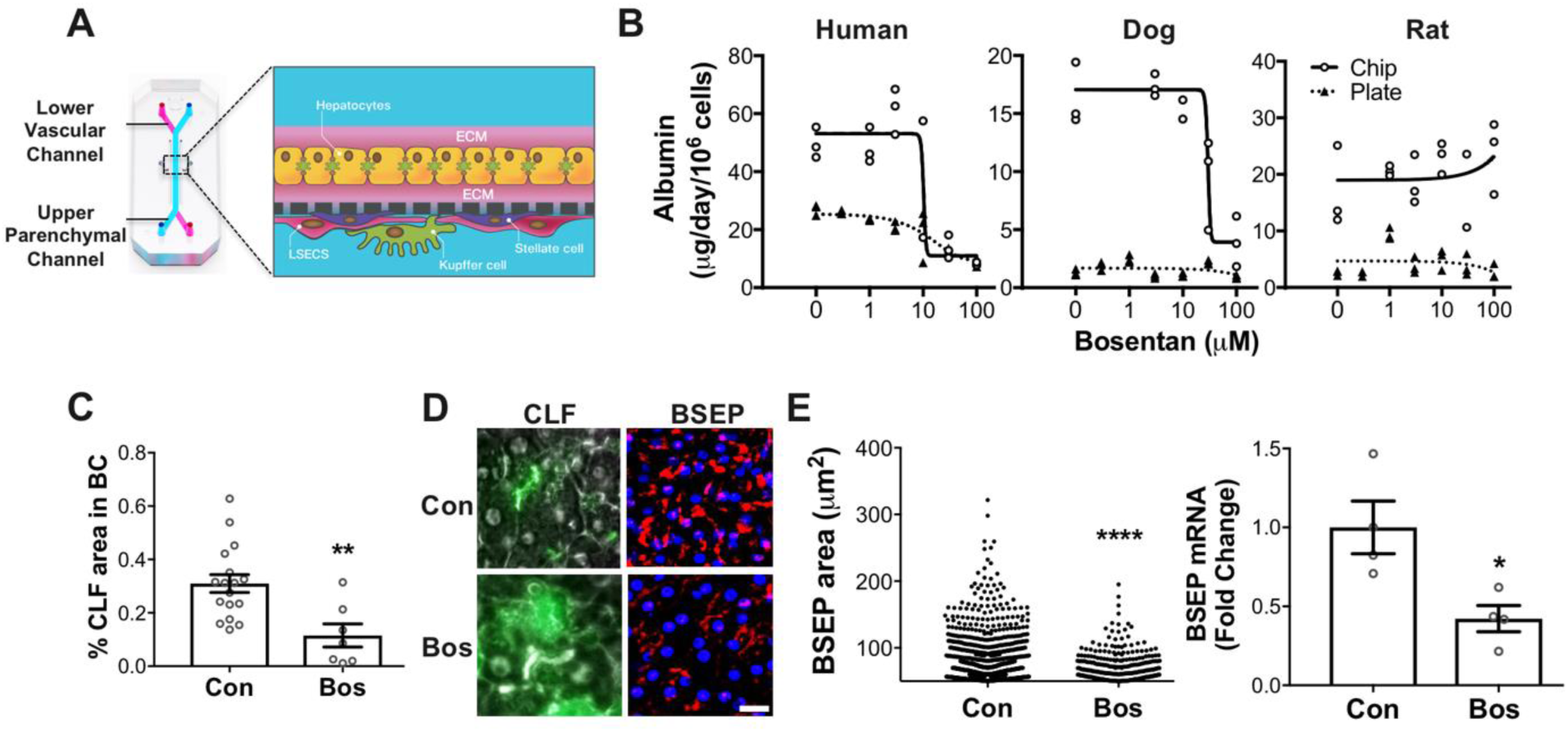
Recapitulation of species-specific drug toxicities in rat, dog, and human Liver-Chips. (**A**) Schematic of the Liver-Chip that recapitulates complex liver cyotoarchitecture. Primary hepatocytes in the upper parenchymal channel in ECM sandwich format and NPCs (LSECs, Kupffer, and stellate cells) on the opposite side of the same membrane in the lower vascular channel. (**B**) Albumin secretion after daily administration of bosentan at 1, 3, 10, 30, or 100 μM for 3 days in dual-cell (hepatocyte and LSECs) human Liver-Chips and plates (hepatocyte sandwich monoculture) and for 7 days in dual-cell dog and rat Liver-Chips and plates (n=3 independent chips and plate wells). (**C**) Quantification of % CLF-positive area in bile canaliculi (BC) from the parenchymal channel after bosentan treatment at 30 μM for 7 days in human Liver-Chips. Mann–Whitney U test (n=3 independent chip with 3 randomly selected different areas per chip, detailed description on the analysis in supplementary materials). (**D**) Representative images of CLF (green, BSEP substrate) and BSEP (red, DAPI in blue) from the parenchymal channel. (**E**) Quantification of BSEP-positive area and fold change of BSEP gene expression. Mann–Whitney U test (n=1 chip for BSEP imaging and n=4 independent chips for gene expression). Scale bar, 20 µm. *P < 0.05, **P < 0.01, ****P < 0.0001. Error bars present mean±SEM.

To assess the liver-specific physiological function of the dual-cell Liver-Chips, we measured secretion of albumin and compared this to results obtained from the same human, rat, and dog hepatocytes cultured alone in the static sandwich culture plates. These studies revealed that all three species-specific Liver-Chips maintained significantly higher (3- to 14-fold greater) levels of albumin production than cells in conventional sandwich monoculture plates (fig. S1C). Importantly, the quantitative range of albumin production we measured in the human Liver-Chip of ~20-70 µg/day/million cells (between days 7 and 14) was very similar to that estimated for humans *in vivo* (50 µg/day/million cells) using *in vitro*-to-*in vivo* extrapolation (iViVE) techniques (17). In contrast, hepatocytes within conventional sandwich monoculture plates showed significantly lower (2.8- to 3.9-fold lower) levels of albumin production than cells in the human Liver-Chip over the same time period.

To further evaluate the physiological relevance of the dual-cell Liver-Chips, the drug metabolizing capacity of the hepatocytes were characterized over time in culture. We measured the activities of the 3 major cytochrome P450 (CYP) isoforms (CYP1A, CYP2B, and CYP3A) that represent key CYP families involved in drug metabolism. We used prototypical probe substrates in a cocktail approach using concentrations of their respective substrates (phenacetin, bupropion, and midazolam) or a single substrate (cyclophosphamide for CYP2B and testosterone for CYP3A for human model) that mirror their Michaelis constant (K_m_) in humans (18). These three isoforms also represent the major CYPs regulated by the xenosensors: aryl hydrocarbon receptor (AhR), constitutive androstane receptor (CAR), and pregnane X receptor (PXR) (19). Importantly, these studies revealed that CYP activities measured in the dual-cell human, rat, and dog Liver-Chips over the 14-day culture period were comparable to, or in some cases greater than, those exhibited by freshly isolated hepatocytes (fig. S2), which are the gold-standard model currently used by pharmaceutical researchers. In contrast, there was a significant decline in CYP activities in all three species in sandwich monoculture plates over the same time period (fig. S2).

## Recapitulation of species-specific drug toxicities in rat, dog, and human Liver-Chips

To explore whether these dual-cell Liver-Chips could be used to predict species-specific drug induced liver injury (DILI) responses, we used the three species models to evaluate hepatotoxic effects induced by bosentan, a dual endothelin receptor antagonist that causes cholestasis in humans, but not in rats or dogs, by inhibiting the bile salt export pump (BSEP) and inducing hepatocellular accumulation of bile salts (20, 21). Daily administration of bosentan at 1, 3, 10, 30, or 100 μM resulted in decreases in albumin secretion with different potencies in these species-specific Liver-Chips, with an IC_50_ of 10, 30 and >100 μM in human, dog, and rat chips, respectively (Fig. 1B). We observed a correlation between the effect in the human Liver-Chip with the clinical response. Notably the concentration in the human Liver-Chip at which we observed toxicity approximated to the plasma concentrations of bosentan (C_max_ = 7.4 μM) that have been associated with DILI in humans (22). In addition, the model was more sensitive in detecting bosentan toxicity compared to sandwich monoculture plates (Fig. 1B) which failed to demonstrate an *in vivo* relevant toxic response. In other complex cell-based liver models including 3D human spheroid hepatic cultures the IC_50_ was found to be more than 10-fold higher than *in vivo* (23). Co-treatment of bosentan (30 μM) with cholyl-lysyl-fluorescein (CLF), a BSEP substrate, inhibited efflux of CLF by 50% (Fig. 1C) resulting in its intracellular accumulation (Fig. 1D) in the Liver-chip consistent with the known mechanism for hepatotoxicity of bosentan in humans. Inhibition of BSEP activity was also accompanied by decreases in BSEP protein and mRNA levels (Fig. 1E). Thus, in addition to recapitulating species-specific hepatotoxicities, these results illustrate that mechanisms of DILI which involve hepatic transporters can be studied in the Liver-Chip. This highlights the advantage of the Liver-Chip in integrating mechanisms of action (in this case BSEP transporter inhibition) to functional outcome (decrease in albumin synthesis) in the same model.

## Detection of diverse phenotypes of hepatotoxicity using quadruple-cell Liver-Chips

To add a higher order of biological complexity to the Liver-Chip necessary to study diverse phenotypes of liver toxicity, we integrated species-specific, non-parenchymal cells (NPC), hepatic stellate and Kupffer cells into the vascular channel to develop the quadruple-cell Liver-Chip model (Fig. 1A). These species-specific, quadruple-cell Liver-Chips also exhibited high levels of albumin secretion similar to those observed in the dual-cell Liver-Chips (fig. S3A). The human and rat quadruple-cell Liver-Chips also maintained high levels of CYP enzyme activities that were similar to, or higher than, those observed in freshly isolated hepatocytes or in the dual-cell Chips (fig. S3B) over long-term culture. We used tool compounds that are known to cause diverse phenotypes of DILI or transaminitis in humans to characterize the ability to use the Liver-Chip for safety and risk assessment in humans and to enable insights into mechanism of action driving the toxicity.

The generic analgesic acetaminophen (APAP) can produce DILI resulting in whole organ failure and death when over-dosed. APAP toxicity is caused by direct injury to hepatocytes mediated by the toxic and reactive metabolite N-acetyl-p-benzoquinone imine (NAPQI) that depletes cellular glutathione (GSH) causing oxidative stress; it can also be detoxified by hepatocytes resulting in formation of glucuronide and sulfate metabolites (24). To evaluate APAP toxicity in the human quadruple-cell Liver-Chip, we maintained a constant flow rate that was determined to best reproduce its metabolism rate and turnover (10 μL/hr of flow rate) based on its known intrinsic clearance. Metabolism of APAP on the Liver-Chip was confirmed by detection of significant amounts of APAP glucuronide in both the parenchymal and vascular channels following daily administration of 3 mM APAP for 20 days (fig. S4A), which confirmed that all four cell types were exposed to the hepatocyte-derived metabolites as a result of diffusion through the porous membrane. Importantly, treatment with APAP resulted in dose-dependent depletion of total GSH and ATP at all concentrations tested (0.5, 3, and 10 mM) in the hepatocytes within the parenchymal channel, and even more potently in the NPCs in the vascular channel (Fig. 2A), highlighting that APAP toxicity is not limited to liver epithelial cells. The depletion of GSH also is suggestive of formation of reactive oxygen species (ROS), which we confirmed by measuring their levels using a fluorescent reporter assay (Fig. 2B). APAP-induced depletion of GSH and ATP was followed by a decline in hepatocyte morphology (fig. S4B) and function, as measured by decreased albumin synthesis and oxidative stress-related injury markers such as alpha glutathione S-transferase (α-GST) and microRNA 122 (miR122) (Fig. 2C). In addition, co-treatment of APAP (3 mM) with the glutathione depleting agent buthionine sulfoximine (BSO; 200 μM) amplified sensitivity to APAP toxicity. This was seen by the increased release of ROS (Fig. 2B), miR122, and α-GST (Fig. 2C) that were not detected at the same APAP concentration in the absence of BSO. Taken together this confirms the reported role of ROS in APAP-induced hepatotoxicity (25).

**Fig. 2.**
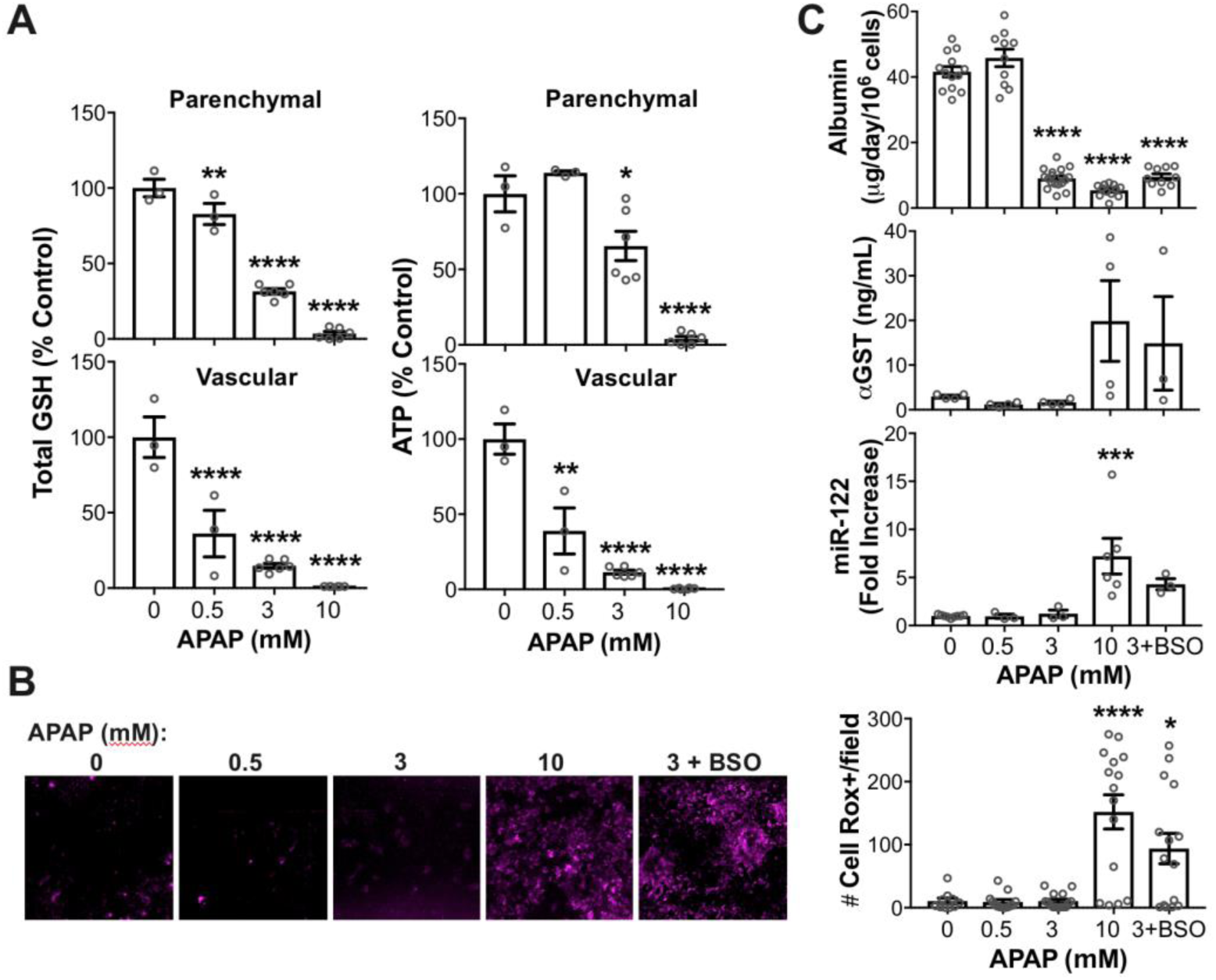
Detection of liver injury and release of various DILI biomarkers using quadruple-cell human Liver-Chips. (**A**) Total GSH and ATP levels from the parenchymal and vascular channels after daily administration of APAP at 0.5, 3, or 10 mM for 7 days in human Liver-Chips. (**B**) Representative images of ROS levels (magenta, CellROX) after daily administration of APAP at 0.5, 3, or 10 mM and co-administration of 3 mM of APAP and 200 μM of BSO for 7 days in human Liver-Chips and quantification of number of CellROX-positive events per field of view. Kruskal-Wallis tests (n=3 independent chip with 3~5 randomly selected different areas per chip). Scale bar, 100 µm. (**C**) Albumin, αGST, and miR-122 secretions from the parenchymal channel after APAP treatment for 7 days in human Liver-Chips. Dunnett’s multiple comparisons test (n=10~18 independent chips for albumin, n=3~9 independent chips for the rest). *P < 0.05, **P < 0.01, ***P < 0.001, ****P < 0.0001. Error bars present mean±SEM.

To explore whether the quadruple-cell Liver-Chips could be used to model DILI mechanisms that target Kupffer cells, we studied JNJ-1, a colony stimulating factor-1 (CSF-1) receptor kinase inhibitor. This compound caused minimal elevations in transaminases (<3-fold) in a human Phase I clinical trial that was attributed to Kupffer cell depletion which can play a role in the clearance of transaminases (26). Very high transaminase levels were reported in two individuals (>10-fold) and were considered idiopathic and unique to the compound (Janssen unpublished data). Minimal dose-related elevations in transaminases (<3-fold) were also observed in rat and dog studies (Janssen unpublished data), but without any correlative microscopic changes in the liver. Kupffer cell depletion observed in the clinic and detected by decreased number of CD68-positive cells was reproduced in human Liver-Chip following administration of JNJ-1 at 3 μM (Fig. 3A), a dose relevant to clinical human concentrations (C_max_ ~2 μM). Kupffer cell depletion was associated further with a decrease in interleukin 6 (IL-6) and monocyte chemoattractant protein-1 (MCP-1) (Fig. 3B) in the vascular channel but was without an effect on hepatocyte function. Importantly, decreased hepatocyte function as measured by lowering of albumin secretion was only observed at 30 μM of JNJ-1, about 15-fold above the human C_max_ (Fig. 3C), suggesting potential for intrinsic toxicity only at high concentrations. These results demonstrated the ability of the human quadruple-cell Liver-Chip to detect a clinically-relevant mechanism of action that targets Kupffer cells, independently of hepatocytes.

**Fig. 3.**
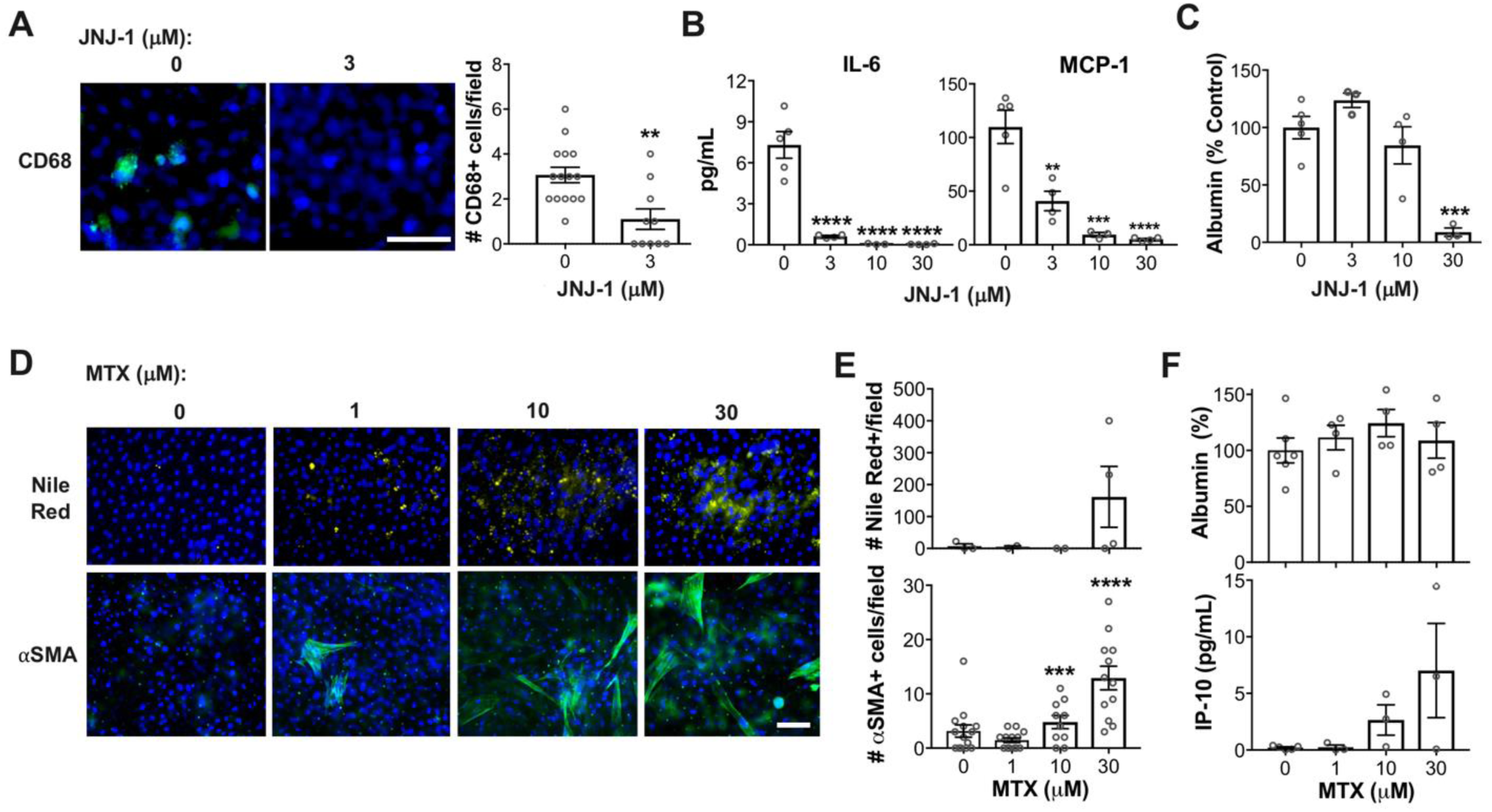
Detection of Kupffer cell depletion, steatosis, and fibrosis in human Liver-Chips. (**A**) Representative images of CD68 (green, DAPI in blue) from the vascular channel after daily administration of JNJ-1 at 3 μM for 7 days in human Liver-Chips and quantification of the number of CD68-positive cells per field of view. Mann–Whitney U test (n=3 independent chip with 3~5 randomly selected different areas per chip). (**B**) IL-6 and MCP-1 release after daily administration of JNJ-1 at 3, 10, or 30 μM for 3 days from the vascular channel in human Liver-chips. (**C**) Albumin secretion after 3 days of JNJ-1 treatment from the parenchymal channel in human Liver-Chips. Dunnett’s multiple comparisons test (n=3~5 independent chips). (**D**) Representative images of lipid droplets (yellow, Nile red, and DAPI in blue) from the parenchymal channel and αSMA (green) from the vascular channel to indicate activated stellate cells after daily administration of MTX at 1, 10, or 30 μM for 7 days in human Liver-Chips. (**E**) Quantification of Nile red-positive events per field of view and αSMA-positive cells per field of view. Kruskal-Wallis tests (n=3 independent chip with 3~5 randomly selected different areas per chip). (**F**) Albumin secretion from the parenchymal channel and IP-10 secretion from the vascular channel after MTX treatment for 7 days and 1 day respectively in human Liver-Chips. Scale bar, 100 µm. *P < 0.05, **P < 0.01, ***P < 0.001, ****P < 0.0001. Error bars present mean±SEM.

## Modeling steatosis and markers of fibrosis on-Chip

Methotrexate (MTX) causes liver injury in humans characterized by steatosis, stellate cell hypertrophy, and fibrosis at maximal plasma concentrations of ~1 μM in some patient populations (27). Importantly, these findings were recapitulated in the human quadruple-cell Liver-Chip where daily administration of MTX at 1, 10, or 30 μM for 7 days resulted in microscopic evidence of lipid accumulation as detected by Nile red staining, and stellate cell activation as indicated by increased expression of α-smooth muscle actin (α-SMA) (Fig. 3D, E). These changes were also associated with increases in interferon γ-induced protein 10 kDa (IP-10), a chemokine whose elevation is associated with liver inflammation and fibrosis (28). There were no abnormalities in albumin secretion (Fig. 3F) or hepatocyte morphology (not shown), which is consistent with the lack of predictive or diagnostic biomarkers for monitoring these toxicities in humans. These studies suggest that inclusion of microscopic endpoints for steatosis and markers of fibrosis in the quadruple-cell Liver-Chip could be an approach to identify compounds with a potential risk for these toxicities.

To investigate whether cross-species Liver-Chip models could be used to predict human-specific steatosis, fialuridine (FIAU) was tested in rat and human quadruple-cell Liver-Chips. Development of FIAU, an anti-viral nucleoside analog, was discontinued in Phase II clinical trials due to liver failure and deaths in 5 out of 15 patients, caused by microvesicular steatosis (29). A review of the animal toxicology data concluded that the studies could not have predicted severe liver injury in humans caused by FIAU (27). Daily administration of FIAU at 1, 10, or 30 µM for 10 days in the human Liver-Chip resulted in a dose-dependent increase in lipid accumulation (Fig. 4A). There was also a concomitant dose-dependent decline in albumin secretion at concentrations ≥ 1 μM, and release of liver injury markers including miR122, α-GST, and keratin 18 (K-18) (Fig. 4B, C). In contrast, there were no effects on lipid accumulation or hepatocyte function following treatment in the rat Liver-Chip with FIAU at the same concentrations and treatment duration as the human Liver-Chip (Fig. 4A, B), which is consistent with past nonclinical studies and species differences observed *in vivo* (29).

**Fig. 4.**
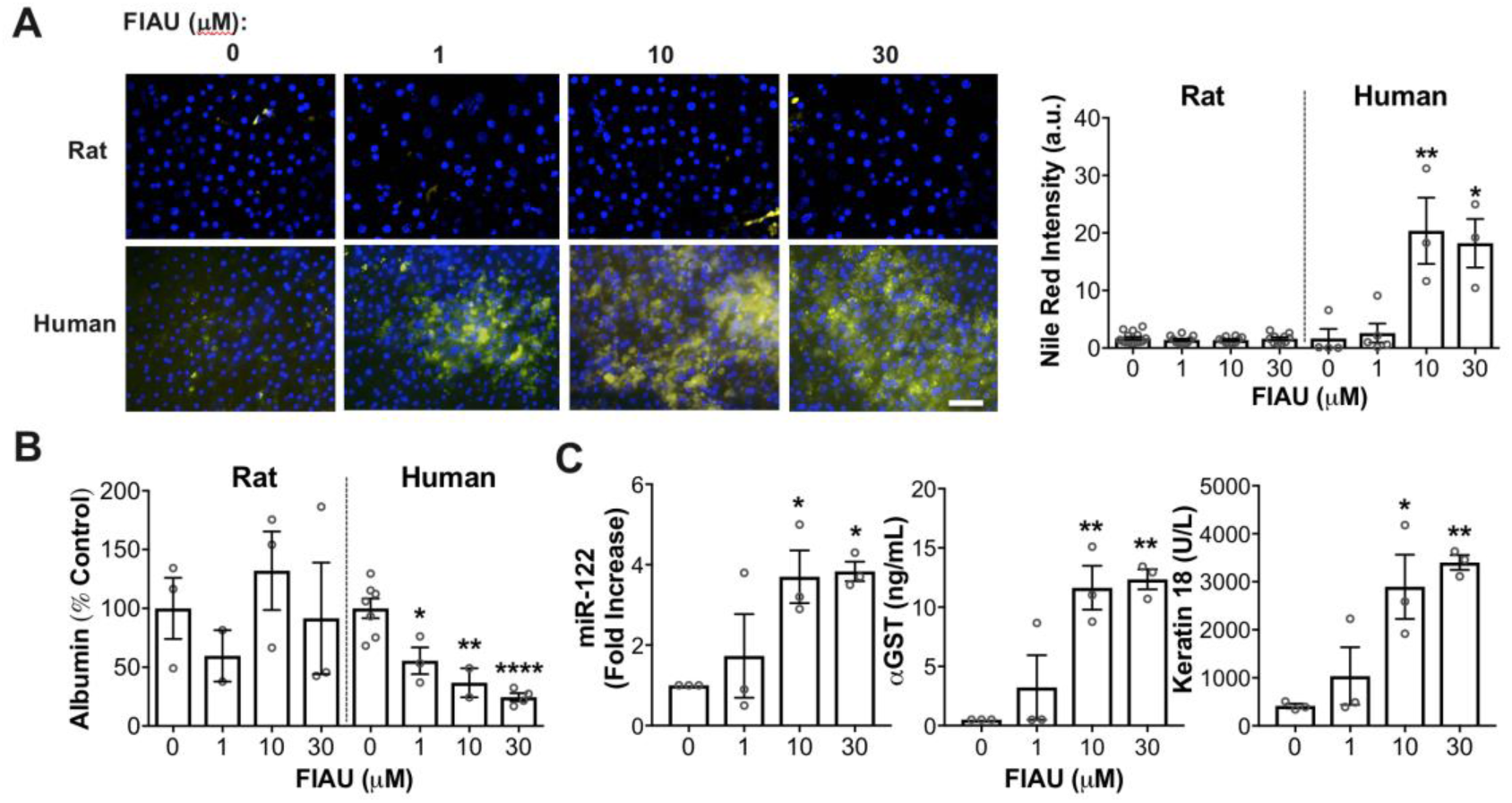
Comparison of species differences in steatosis using rat and human Liver-Chips following fialuridine (FIAU) treatment. (**A**) Representative images of lipid droplets (yellow, Nile red, and DAPI in blue) from the parenchymal channel after daily administration of FIAU at 1, 10, or 30 μM for 10 days in rat and human Liver-Chips and quantification of Nile red intensity. (**B**) Albumin secretion as % control after FIAU treatment for 7 days in rat and human Liver-Chips. (**C**) Mir-122, αGST, and keratin 18 secretions after FIAU treatment for 10 days in human Liver-Chips. Dunnett’s multiple comparisons test (n=3 independent chips). Scale bar, 100 µm. *P < 0.05, **P < 0.01, ****P < 0.0001. Error bars present mean±SEM.

## Use of species-specific Liver-Chips to query human relevance of animal liver toxicities

It is not uncommon for compounds to be discontinued due to liver toxicity observed in rats or dogs prior to testing in humans because of uncertainties on the human relevance of these findings. To evaluate whether species-specific Liver-Chips could be used to assess human relevance, a Janssen proprietary compound that was discontinued due to liver toxicity in rats was characterized in the cross-species Liver-Chips. Daily oral administration of JNJ-2 to rats for 2 weeks resulted in liver fibrosis, supported by increased α-SMA staining within stellate cells, at 150 mg/kg (C_max_ = 11.6 μg/mL; 24 μM) that was persistent 3 months after compound wash-out (Fig. 5A). These findings were associated with chronic inflammation of portal areas and decreases in albumin with no changes in transaminases in rats, and as a result, JNJ-2 was discontinued prior to testing in non-rodent species. Interestingly, daily treatment of JNJ-2 at 3, 10, or 30 µM in rat quadruple-cell Liver-Chip for 4 days resulted in a dose-dependent increase in expression of α-SMA, a marker for fibrosis, within stellate cells (Fig. 5B, C); decreases in albumin were observed, but did not reach significance (Fig. 5D). In contrast, treatment of human Liver-Chips at the same concentrations did not produce these abnormalities, even when extended for 14 days of treatment (Fig. 5B, C, D). These results imply a potential species differences in response between rats and humans with JNJ-2.

**Fig. 5.**
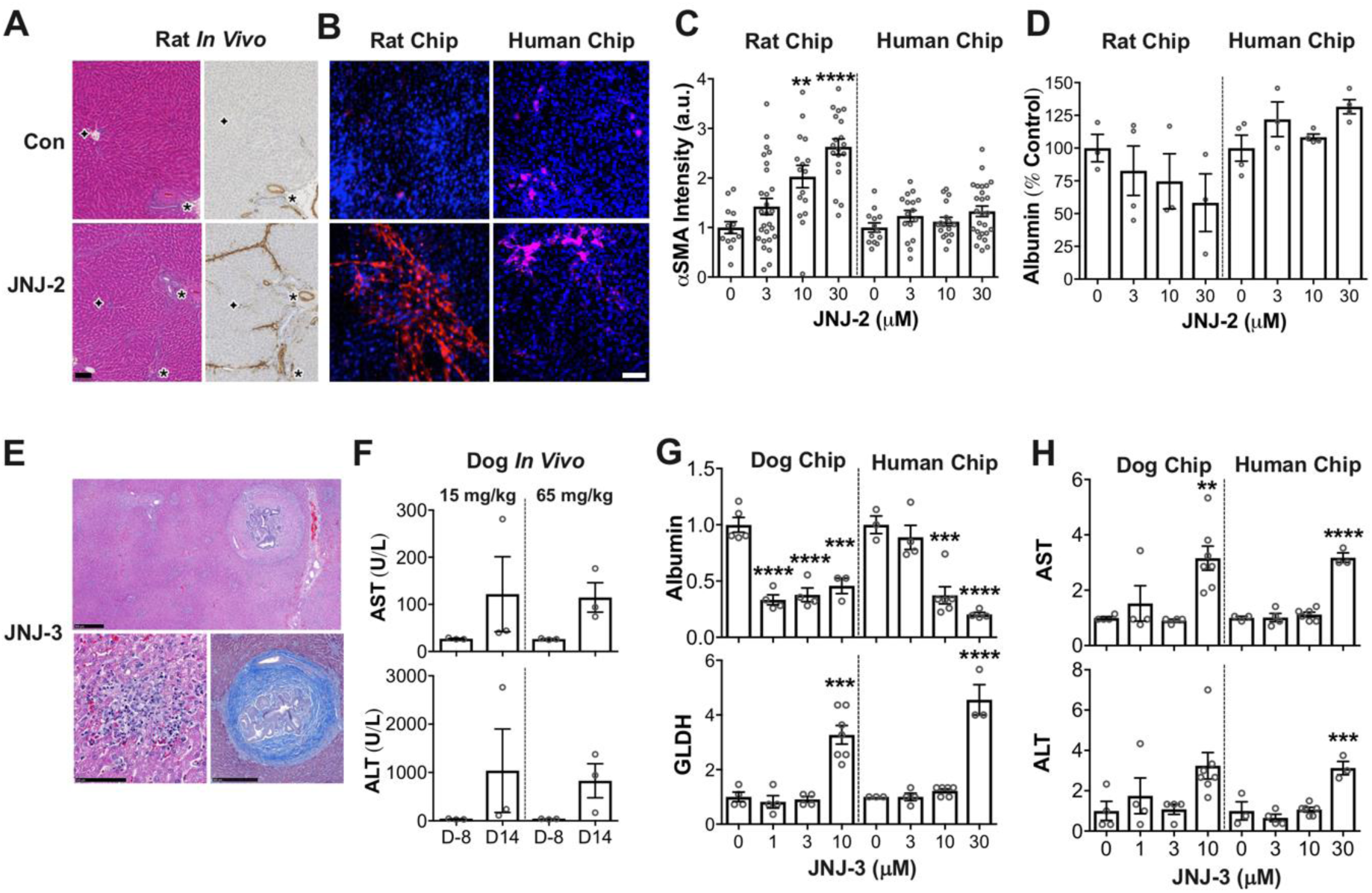
Comparison of species differences in fibrosis using rat and human Liver-Chips and elevation of transaminases using dog and human Liver-Chips. (**A**) H&E and αSMA images after daily oral administration of JNJ-2 at 150 mg/kg for 2 weeks in rats. (**B**) Representative images of αSMA (red, DAPI in blue) from the vascular channel after daily administration of JNJ-2 at 30 μM for 4 days in rat Liver-Chips and 14 days in human Liver-Chips. (**C**) Quantification of αSMA intensity from the vascular channel after daily administration of JNJ-2 at 3, 10, or 30 μM for 4 days in rat Liver-Chips and 14 days in human Liver-Chips. (**D**) Albumin secretion as % control after daily administration of JNJ-2 for 4 days in rat Liver-Chips and 14 days in human Liver-Chips. (**E**) Liver changes (H&E and Masson’s Trichrome staining) in the dogs following daily administration of JNJ-3 at 65 mg/kg for 14 days. Low magnification showing multifocal areas of necrosis, inflammation and portal fibrosis in the hepatic parenchyma. High magnification showing necrotic hepatocytes with inflammatory cells (left) and moderate portal fibrosis (collagen stained as blue), inflammation, and bile duct hyperplasia. (**F**) AST and ALT measurements after JNJ-3 treatment at 15 and 65 mg/kg/day in dogs for 6 days. (**G**) Fold increases of albumin and GLDH secretion from the parenchymal channel after administration of JNJ-3 at 1, 3, or 10 μM for 1 day in dog and human Liver-Chips. (**H**) Fold increases of AST and ALT secretion from the parenchymal channel after administration of JNJ-3 for 1 day in dog and human Liver-Chips. Dunnett’s multiple comparisons test (n=3~4 independent chips). Scale bar, 100 µm. *P < 0.05, **P < 0.01, ****P < 0.0001. Error bars present mean±SEM.

We also tested another Janssen proprietary compound, JNJ-3, that was discontinued from further development due to hepatocellular necrosis, portal fibrosis, and biliary hyperplasia following daily dosing at 15 and 65 mg/kg for 14 days in dogs at maximal plasma concentrations of 12.7 and 19.4 μM, respectively (Fig. 5E). These findings were associated with significant elevations of ALT and AST after 6 or 14 days of treatment (Fig. 5F). JNJ-3 was administered to dog and human Liver chips for 4 days after which the study was stopped due to significant damage to hepatocytes at the highest concentrations tested, and biochemical endpoints were measured in samples collected at 24 hours post-dose. Administration of JNJ-3 significantly decreased albumin secretion at ≥ 1 μM in dog Liver-Chips, and at ≥10 μM in human Liver-Chips (Fig. 5G). ALT, AST, and glutamate dehydrogenase (GLDH) were also elevated in dog and human Liver-Chips, but only at the highest concentrations of 10 and 30 μM, respectively (Fig. 5G, H), indicating that for this compound albumin is a more sensitive marker of hepatocyte dysfunction in the Liver-Chip model. Transaminase elevations in dog Liver-Chip occurred at concentrations that bridged dog plasma concentrations. Thus, the dog Liver-Chip corroborates *in vivo* results. Although toxicity by JNJ-3 was 3 to 10-fold more potent in dog Liver-Chip than human chip, it is highly probable that liver toxicity could have occurred in patients if this compound had progressed into clinical trials.

## Identifying risk for idiosyncratic DILI using the Human Liver-Chip

One of the most difficult forms of hepatotoxicity to predict in the clinic relates to idiosyncratic DILI responses that are often missed during nonclinical and early clinical testing. To explore whether the human Liver-Chip might be useful to predict these types of response, we tested TAK-875, a GPR40 agonist that was discontinued in Phase III trials due to low incidence (2.7%) treatment-related elevations in transaminases (>3-fold rise in upper limit of normal) combined with a few individual cases of serious DILI (30). *In vitro* and *in vivo* studies identified formation of reactive acyl glucuronide metabolites, suppression of mitochondrial respiration, and inhibition of hepatic transporters by TAK-875 as potential mediators of its hepatotoxic effects (31). Daily administration of TAK-875 at 10 μM (equivalent to human C_max_) in the human quadruple Liver-Chip for approximately 2 weeks resulted in formation of the oxidative metabolite (M-1) (6.4% of parent) (Fig. 6A) at similar levels to that reported in humans (10% of parent) (32), and production of significant amounts of the acyl glucuronide metabolite (54% of parent) (TAK-875AG), again consistent with reports that glucuronidation of TAK-875 represented a major clearance pathway in humans. Glucuronide metabolites are substrates for canalicular and basolateral hepatic MRP transporters, but at high intracellular concentrations they may inhibit their own efflux and accumulate in hepatocytes. Daily administration of TAK-875 at 3, 10, or 30 μM in Liver-Chip resulted in a dose-dependent decrease in biliary efflux of the MRP2 substrate 5(6)-Carboxy-2′,7′-dichlorofluorescein diacetate (CDFDA), implying MRP2 inhibition by TAK-875AG formed in the human Liver-Chip (Fig. 6B). This allowed us to probe the consequences of prolonged exposures to TAK-875 and its formed reactive metabolite TAK-875AG in Liver-Chip in a 2-week study. We found an effect on mitochondrial membrane potential, confirmed by a dose-related and time-dependent redistribution of the mitochondrial potential sensitive dye tetramethylrhodamine methyl ester (TMRM) detected at 1 week of treatment (Fig. 6B,C), and lipid droplet accumulation and formation of ROS (Fig. 6B,C) at the end of the 2-week treatment; these endpoints may be consequential to perturbation of the mitochondria. We then investigated whether an innate response could be detected in Liver-Chip treated with TAK-875 based on the prevailing hypothesis for immune-mediated DILI that an innate response caused by covalent protein binding, cell stress, and release of damage associated molecular patterns (DAMPs) can trigger an adaptive immune attack of hepatocytes in a few susceptible individuals (33). Treatment with TAK-875 caused a biphasic and significant release of the inflammatory cytokines MCP-1 and IL-6 at 10 µM (human equivalent C_max_) but not at 30 µM (Fig. 6D); lack of cytokine release at higher concentrations could be due to reduced cell health resulting from ROS formation.

**Fig. 6.**
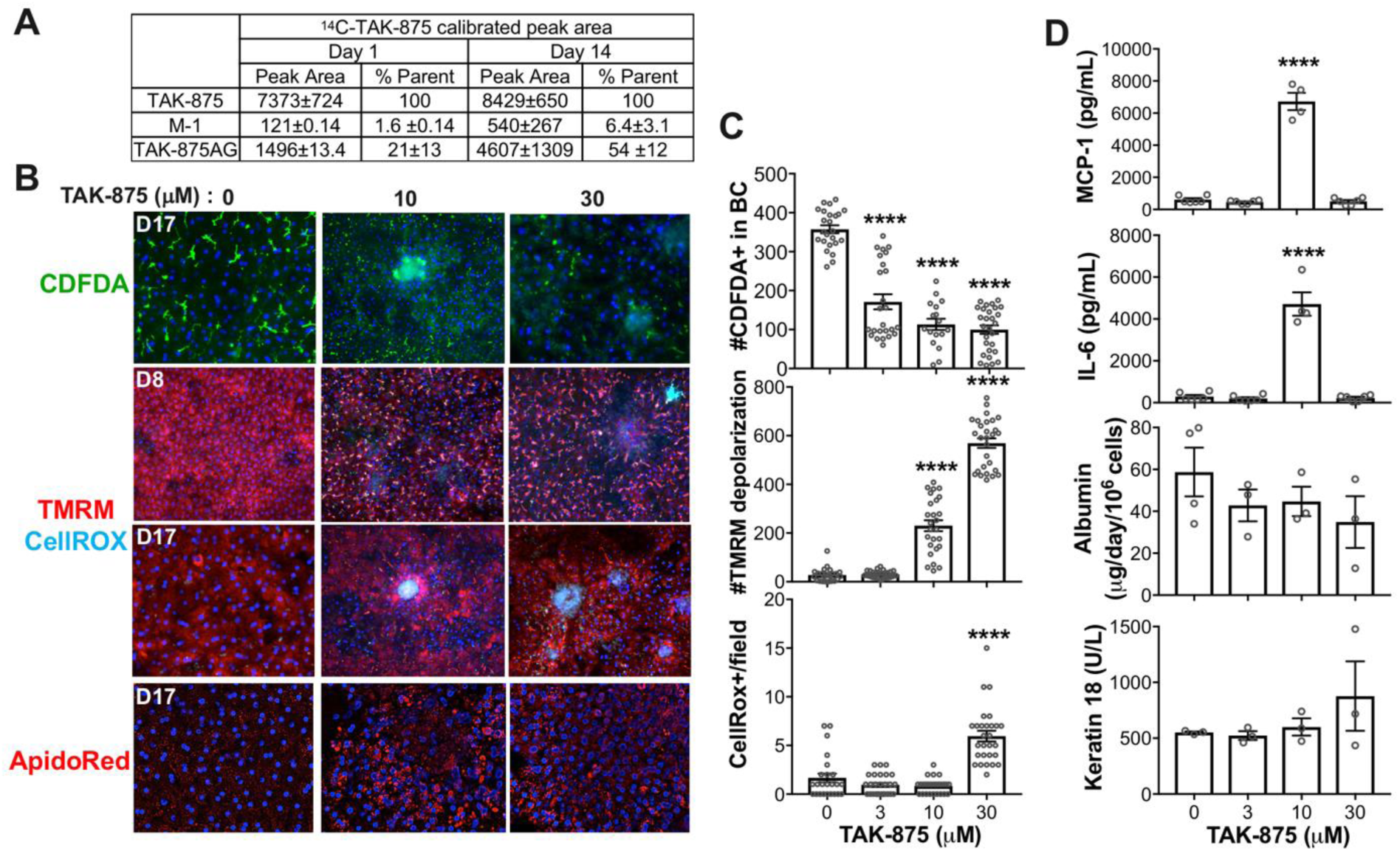
Identifying risk for idiosyncratic DILI using Human Liver-Chip. (**A**) Formation of TAK-875 metabolites after day 1 and day 14 of TAK-875 treatments in human Liver-Chips. (**B**) Representative images of CDFDA (green, DAPI in blue) to identify MRP2 transporter activity, TMRM, and CellROX (red and cyan respectively, DAPI in blue) to detect mitochondrial depolarization and ROS respectively, and AdipoRed (red, DAPI in blue) to detect lipid droplets after daily administration of TAK-875 at 10 or 30 μM for 8 days or 15 days in human Liver-Chips. (**C**) Quantifications of number of CDFDA positive fractions in bile canaliculi area, number of redistributed TMRM fractions, and CellROX positive events per field of view after daily administration of TAK-875 at 3, 10, or 30 μM for 15 days in human Liver-Chips. Kruskal-Wallis tests (n=3 independent chip with 5 randomly selected different areas per chip). (**D**) MCP-1 and IL-6 releases from the vascular channel and albumin and keratin 18 secretion from the parenchymal channel after 14 days of TAK-875 treatment in human Liver-Chips. Dunnett’s multiple comparisons test (n=3 independent chips). ****P < 0.0001. Error bars present mean±SEM.

## Discussion

These results suggest that species-specific Liver-Chips can be used to predict rat, dog, and human hepatotoxicities, and that they may be used to estimate human relevance of liver toxicities seen in animals for safety and risk assessment. They also could potentially be used to predict human idiosyncratic hepatotoxicities and elucidate mechanisms of action.

Ongoing research on DILI ranges from basic understanding of adverse pathways and mechanisms by which some compounds cause DILI to clinical research aimed at identifying predictive DILI biomarkers. This research is spurred by human safety concerns for liver failure and the associated high costs incurred when these drugs are withdrawn from clinical trials or from the market based on incorrect predictions from nonclinical testing. Recently, the FDA released a Predictive Toxicology Roadmap aimed at qualifying *in silico* and *in vitro* models to enable their use in regulatory decision making; qualification of these models will encourage their uptake and use, which would also meet objectives of reduction, refinement, and replacement of animals. While there are significant advances in the development of *in vitro* models that predict DILI, key gaps remain that need to be addressed. For example, lack of metabolic competence in plated primary hepatocytes has been addressed by more complex 2D co-culture or 3D spheroid models, which can maintain metabolic competence for prolonged durations (14, 34–36). But these models are static and closed, therefore, metabolites may accumulate to levels that are not physiologically relevant or they may be underrepresented due to loss following medium replenishment that is required to maintain cell survival. In addition, dense 3D constructs often suffer from limited oxygen and nutrient transport, and even when spheroid cultures include endothelial cells and Kupffer cells, they lack physiologically relevant tissue-tissue interfaces and cytoarchitecture that has been demonstrated to be a key driver of cell function (10, 37). It is also impossible to apply physiologically relevant fluid flow and associated mechanical forces in which drugs pass specifically through an endothelium-lined interface as they do *in vivo*. Mechanical forces have also been demonstrated to be a key player in gene expression and *in vivo* cell function (10). In addition, in spheroids it is difficult to visualize dynamic changes in cell position, morphology, or function within these large, disorganized, and multi-cellular structures.

Recently, there has been many attempts to develop liver-based cell systems to replicate organ-level functions using microengineered approaches, such as microphysiological systems (MPS), and some have included the four cell types we included in the quadruple-cell Liver-Chips in the present study (38–40). Some of these studies demonstrated enhanced functionalities with flow (e.g., increased albumin secretion and CYP activities) but were limited to short-term (e.g., 1 day) cultures (38). In contrast, in the present study, we showed that both the dual-cell and quadruple-cell Liver-Chips remain metabolically competent and maintain albumin production as well as activities of multiple key drug metabolizing enzymes at *in vivo*-like levels for at least 14 days in culture. More importantly, while DILI responses to various drugs were measured in other MPS liver models (36, 39, 41), the drug concentrations utilized were not clinically relevant. In contrast, we used drug levels similar to those observed in the plasma of animals and patients in the present study, and as a result, we were able to generate results that closely recapitulate those previously reported in both nonclinical animal studies and human clinical trials.

Endpoints assessed in most *in vitro* systems are limited to measures of cell viability as an initial assessment of potential hazards, but they often do not capture the mechanisms that underlie DILI or clinical relevant endpoints, and they are not effective for human risk assessment. We evaluated whether the Liver-Chip could detect more complex and mechanistically relevant DILI endpoints. Using a combination of microscopy, tissue staining, and measurement of DILI biomarkers, we were able to detect diverse phenotypes of DILI including hepatocellular injury, cholestasis, steatosis, Kupffer cell depletion, and stellate activation as a marker of fibrosis. Thus, the quadruple-cell Liver-Chip appears to be suitable for detecting toxicities that are attributable to direct effects on the four liver cell types included in our model; however, they are currently not capable of detecting toxicities of the bile duct as this cell type has not yet been included in the design of the current Liver-Chip. It was interesting to note that, with the tool compounds tested so far, toxicities in the model were detected at concentrations that bridged human plasma levels associated with DILI, suggesting that the model has potential to be used for human risk assessment.

Although the enzymatic activities in human Liver-Chip were robust, some activities were lower or higher in comparison to fresh human hepatocytes, which most likely reflects donor to donor variability. The CYP enzyme activities in rat Liver-Chip were similar to those measured in fresh rat hepatocytes, when a single substrate was used, but they were higher when a cocktail of CYP substrates were used, which suggests potential for interactions among these substrates in the cocktail approach (fig. S2, S3B).

One advantage of the microengineered Liver-Chips is the use of continuous flow in an open system, which ensures that all cells are exposed sufficient levels of the parent drug and its metabolites simply by adjusting the flow rate. The open system also allows for continuous collection or sampling of the effluents of both the vascular and parenchymal channels, which prevents accumulation of the parent and its metabolites, while enabling measurement of biomarkers and other biology endpoints over time.

The ability to measure mechanistic endpoints and biomarkers in the model also makes it suitable for delineating pathways and mechanisms causing DILI. For instance, the observation that GSH and ATP depletion are early events in APAP-mediated toxicity followed by a decline in hepatocyte function and finally by oxidative stress and overt injury, suggests that toxic metabolite-mediated mitochondrial dysfunction and ATP depletion are likely early events in the APAP toxicity cascade. Indeed, mitochondrial dysfunction has been identified as a hazard for APAP toxicity (42). Depletion of GSH and ATP in non-parenchymal cells following treatment with APAP implies that the toxic metabolite can escape hepatocytes and mediate an effect on other cell types. This is supported by studies in mice where APAP metabolites were detected bound to reticulocytes, which lack drug metabolizing enzymes, indicating that these metabolites also escape hepatocytes *in vivo* (43). The increased sensitivity of NPCs to APAP toxicity compared to hepatocytes may be due to a reduced detoxification capacity relative to that in hepatocytes. The current studies in Liver-Chip confirm that both hepatocytes and non-parenchymal cells contribute to APAP hepatoxicity *in vivo*.

Specific contexts of use must be defined for predictive *in vitro* models, such as the Liver-Chip, prior to their qualification to make key decisions in the drug development process. These contexts of use can be developed in areas such as prediction of human liver toxicity, human relevance of toxicity observed in animal studies, or identifying DILI potential of compounds that form reactive metabolites. Results of our studies with bosentan, fialuridine, methotrexate, and JNJ-1 show that the human Liver-Chip can be used to study diverse mechanisms of responses in the liver with compounds that target parenchymal and non-parenchymal cells, and more importantly, that these occur at clinically relevant concentrations that bridge plasma concentrations where the effects were observed in humans. Human-specific sensitivities to toxicity by bosentan and FIAU were also confirmed in the human Liver-Chip compared to the companion animal Liver-Chips demonstrating specie-specific *in vivo*-*in vitro* correlation. The putative mechanism for bosentan — inhibition of bile acid efflux via BSEP resulting in intracellular accumulation of bile acids — was also confirmed; an advantage of the Liver-Chip is that we could couple these mechanistic endpoints to a measurable decline in hepatocyte function. The species differences in bosentan toxicity may be related to species differences in the composition of toxic bile acids (21). The species differences for FIAU is explained by species differences in expression and activity of the nucleoside transporter 1 (EMT1), which is absent in rats, but present in humans where it facilitates entry of FIAU into the mitochondrial membrane causing mitochondrial toxicity (44). Inhibition of CSF-1 receptor kinase by JNJ-1 caused elevation in transaminases in animals and humans that is considered secondary to Kupffer cell depletion. This was confirmed in Liver-Chip where treatment with JNJ-1 caused a depletion in Kupffer cells accompanied by decreases in cytokines. Transaminase elevations have been observed with other agents that inhibit CSF-1/CSF-1R (45). Kupffer cells depend on CSF-1 for viability and evidence has been presented that transaminase elevations are downstream of Kupffer cell depletion and possibly a consequence of a role for Kupffer cells in transaminase clearance (26).

A gap exists in assigning human relevance of liver toxicities observed in animal studies, especially when these are observed in only one species. JNJ-2 and JNJ-3 are examples of compounds that caused liver toxicity in animal studies and were discontinued prior to clinical development; JNJ-2 caused fibrosis in rats, a finding that is not monitorable in humans, while JNJ-3 caused severe hepatocellular injury and biliary hyperplasia in dogs as early as a week after daily treatment. The rat Liver-Chip was very sensitive to treatment with JNJ-2, while no toxicity was observed in the human Liver-Chip at the same concentrations up to 14 days of daily treatment. Activation of stellate cells noted in rat, but not in human Liver-Chip, confirmed that we could reproduce the pathophysiology observed in rats in the rat Liver-Chip. Moreover, lack of a similar response in the human Liver-Chip suggests potential for species difference in the response. Although these results are interesting and could have influenced an internal decision to test the compound in non-rodents to address whether fibrosis was rat-specific, the model would need robust qualification with a specific context of use to convince regulatory agencies to make a decision with regards to the lack of human relevance of the rat findings. As we observed release of cytokines/chemokines including IP-10 in Liver-Chips treated with MTX, FIAU, and JNJ-2 which has been identified as a potential marker for fibrosis (46), this model also may be amenable biomarker discovery, especially for more challenging disease areas, such as steatosis and fibrosis, where suitable biomarkers for monitoring in humans are lacking.

Reactive metabolite formation has been identified as an important hazard associated with compounds that cause rare or idiosyncratic DILI. The microengineered Liver-Chip provides an opportunity to put formation of reactive metabolites in a cell, tissue, and organ context for a functional readout of their contribution to DILI. For instance, putative adverse pathways for TAK-875-mediated DILI were detected in discrete *in vitro* models (30), however, the physiological consequence of perturbing these mechanisms could not be studied. Treatment of Liver-Chip with TAK-875 showed that continuous and prolonged exposure to parent and reactive metabolites caused mitochondrial dysfunction, oxidative stress, formation of lipid droplets, and an innate immune response (i.e., cytokine release), all of which are harbingers of DILI for patients with increased susceptibility. Interestingly, in a 24-week Phase III clinical trial with TAK-875, liver biopsies of 5 of 7 patients that had abnormal transaminase levels in the study also presented with steatosis (47). However, it was challenging to assign causality because of disease background in Type 2 diabetic patients, but a treatment related effect could not be ruled out. The Liver-Chip studies suggest that steatohepatitis secondary to mitochondrial dysfunction and ROS formation could be a potential phenotype of DILI following treatment with TAK-875. This exemplifies the advantage of the Liver-Chip for assessing the pathophysiological consequence of reactive metabolite formation, which has been strongly associated with idiosyncratic DILI (33, 48, 49).

In conclusion, we have shown that species-specific Liver-Chips have potential future application for safety testing, disease modeling, mechanism of action determination, biomarker identification, and predicting human hepatotoxicities, including idiosyncratic responses. This approach also could be used to query human relevance of toxicities observed in nonclinical animal studies or for mechanistic investigations of DILI detected in nonclinical and clinical studies.

## Acknowledgments

We thank Shannon Dallas, Jose Silva, Peggy Guzzie-Peck, and Gilles Bignan for useful scientific discussions, Linda Shanno for gene expression analyses, Jie Chen for bioanalysis of APAP metabolites, Michael Kelley for critical review of the manuscript, Victor Antontsev for iViVE modeling, Utkarsh Doshi at IVAL for performing the LC-MS analysis, and Joshua Resnikoff and Payal Patel for their contribution in initial model validation.

## Funding

Primary funding support for this project was obtained from Janssen Biotech, Inc., a Janssen Pharmaceutical company of Johnson & Johnson, with additional funding from AstraZeneca, the Wyss Institute for Biologically Inspired Engineering at Harvard University, and the Defense Advanced Research Projects Agency (DARPA) under Cooperative Agreement Number W911NF-12-2-0036.

## Author contributions

G.A.H, M.A.O, K-J.J, L.E, and D.E.I initiated the project, design the model features for dual-cell model, defined model development strategy, and provided guidance to solve initial challenges, successful model development and characterization; G.A.H, M.A.O, K-J.J, and D.E.I initiated the project, design the model features for quadruple-cell model, and defined model development and characterization strategy; M.A.O, K-J.J, H-K. L, D.S, and G.A.H designed quadruple-cell model characterization study experiments; L.E, K-J.J, and G.A.H designed dual-cell model characterization experiments which were included in the supplementary figures; M.A.O, D.S, H-K. L, and K-J.J designed all compounds studies; K-J.J, J.R, K.K, D.P, G.K, J.E.R, and D.C performed all Liver-Chip experiments; J.N performed CLF efflux imaging and analysis in bosentan study; R.B performed image analysis of JNJ-2 study; H.P performed dog Kupffer and stellate cell isolation; J.S performed iViVE analysis; H-K. L, J. C and W.L performed met ID analysis of APAP, TAK-875, and compound analysis for PDMS absorption of JNJ compounds; M.S validated and performed liver biomarker analyses; M. D, M.S, T.S, C.M, E.B and B.S designed, conducted or analyzed *in vivo* experiment for JNJ compounds; B.J designed a cocktail substrate for CYP enzyme activity test; A.S and L.A. performed LCMS analysis for CYP450 enzyme activity measurement; A.H and S.H were involved initial model development; J.R, K.K, J.E.R, J.E, M.S, E.B, B.S, K-J.J, H.L, and M.S wrote the methods; K-J.J prepared all figures; K-J.J and M.A.O wrote the rest of manuscript with editing and input from D.E.I, G.A.H, L.E, D.W, E.B, B.S, C.M, H-K.L, J.R, A.H, D.P, J.E.R, R.B, and J.S.

## Competing interests

D.E.I. holds equity interest in Emulate, Inc. and chairs its scientific advisory board. K-J.J, J.R, K.K, D.P, G.K, J.N, D.C, R.B, J.S, and G.A.H are employees of and hold equity interests in Emulate, Inc.

## Data and materials availability

All data are available in the main text or the supplementary materials.

## Materials and Methods

### Cell sourcing

Cryopreserved primary human hepatocytes were purchased from Triangle Research Labs (Lonza, Morrisville, NC, USA) and Gibco (Thermo Fisher Scientific, Waltham, MA, USA); cryopreserved primary rat hepatocytes and freshly isolated dog hepatocytes were purchased from Biopredic (Saint Grégoire, France) and QPS (Newark, DE, USA), respectively. Cryopreserved primary human liver sinusoidal endothelial cells (LSECs) were purchased from Cell Systems (Kirkland, WA, USA), rat and dog LSECs were purchased from Cell Biologics (Chicago, IL, USA) and each were cultured according to their respective vendor protocols. Cryopreserved human and rat Kupffer cells were purchased from Thermo Fisher Scientific, human and rat stellate cells were purchased from Lonza, and each were cultured according to their respective vendor protocols. Dog Kupffer and stellate cells were isolated from hepatic NPC (QPS) at Emulate, Inc. following protocols by Olynyk et al. (1) and Riccalton-Banks et al. (2).

### Liver-Chip culture

Prior to cell seeding, chips (S-1 Chips, Emulate, Inc. Boston, MA, USA) were functionalized using Emulate’s proprietary protocols and reagents (ER™). After surface functionalization, both channels of the Liver-Chip were coated with species-specific extracellular matrices (ECM). For the human Liver-Chip, we used a mixture of collagen type I (Corning, Corning, NY, USA) and fibronectin (Gibco); for the rat Liver-Chip we used a mixture of collagen type IV (Sigma-Aldrich, St. Louis, MO, USA) and fibronectin (Gibco); for the dog Liver-Chip we used a mixture of collagen I (Corning), collagen type IV (Sigma-Aldrich), and fibronectin (Gibco). Primary hepatocytes were seeded in the upper channel of the Liver-Chips at a concentration of 3.5 million cells/mL and later overlaid with Matrigel® (Corning), then incubated at 37°C, 5% CO_2_. For the dual-cell culture model (hepatocytes and LSECs), the LSECs were seeded at a concentration of 2-4 million cells/mL in the lower vascular channel. For the quadruple-cell model (hepatocytes, endothelial cells, Kupffer, and stellate cells), a mixture of LSEC, Kupffer, and stellate cells were seeded in the channel of the Liver-Chips at the following concentrations: 3 million cells/mL for LSEC, 0.5 million cells/mL for Kupffer cells, and 0.1 million cells/mL for stellate cells. After cell seeding, the upper channel of the Liver-Chip was maintained in William’s E Medium (WEM) containing Glutamax (Gibco), ITS+ (Corning), dexamethasone (Sigma-Aldrich), ascorbic acid (Sigma-Aldrich), fetal bovine serum (Sigma-Aldrich), and Penicillin/Streptomycin (Sigma-Aldrich). The vascular channel of the Liver-Chip was maintained with species-specific endothelial media (Emulate, Inc.). Two days after seeding, the Liver-Chips were connected to the Human Emulation System™ (Emulate, Inc.) and both of the chip channels were perfused at 30 µL/hr to provide a continuous supply of fresh media for the duration of the experiments.

### Immunofluorescence staining

Liver-Chips and static sandwich monoculture plates were fixed with 4 % paraformaldehyde for 15 minutes at room temperature, washed with PBS, and permeabilized (1 % saponin and 1 % BSA in PBS). Blocking (10 % serum and 1 % BSA in PBS) and incubation with the primary antibodies in the blocking buffer for overnight at 4°C was followed by a two-hour incubation with secondary antibodies (Cell Signaling, Danvers, MA, USA) in the blocking buffer at room temperature. Immunostaining was performed with specific primary antibodies (anti-MRP2, anti-stabilin-1, anti-BSEP, anti-α-SMA, anti-CD68; Abcam, Cambridge, MA, USA) and images were acquired with either an Olympus fluorescence microscope (IX83) or Zeiss confocal microscope (AxiovertZ1 LSM880).

### Live cell staining

Liver-Chips were stained in the upper channel with 5(6)-Carboxy-2′,7′-dichlorofluorescein diacetate (CDFDA) (Thermo Fisher) to visualize bile canaliculi and MRP2 activity, cholyl-lysyl-fluorescein (CLF) (Corning) to visualize bile canaliculi and BSEP activity, Nile red (Thermo Fisher) or AdipoRed (Lonza) to visualize lipid droplet accumulation, Tetramethylrhodamine, methyl ester (TMRM) (Thermo Fisher) to visualize active mitochondria, and CellROX® (Thermo Fisher) to visualize cellular oxidative stress. Each staining solution was prepared in blank medium or compound dosing medium and added to the upper channel, incubated for 15 minutes at 37 °C, and washed three times with medium. The stained chips were imaged using a specific filter, and were de-blurred with Olympus cellSens software. Using ImageJ-Fiji, the fluorescent images were histogram adjusted to remove background, followed by fluorescence quantification using the Integrated Density calculation function.

### Biochemical assays

Albumin secretion from the upper channel was quantified using ELISA kits for human, rat, and dog models effluent samples (human and rat: Abcam; dog: Immunology Consultants Laboratory, Inc., Portland, OR, USA) and the assays were performed according to the protocol provided by each vendor.

### ATP and GSH

ATP was measured using a modified CellTiter-Glo® Luminescent Cell Viability Assay (Promega, Madison, WI, USA). Briefly 50 μL of cell lysate was mixed with 50 μL of ATP luminescent reaction mixture, incubate for 5 min, and luminescent intensity measured using a luminescence plate reader according to manufacturer’s instruction. ATP concentrations in the lysates was quantified with an ATP standard curve.

Total glutathione was measured by adding, 50 μL of cell lysate from each sample to 50 μL of the GSH-Glo™ Reagent 2X (Promega) with DTT (~2.5 mM final) reagent (to reduce all glutathione) to each well of a 96-well plate. Samples were incubated at room temperature for 30 minutes. Reconstituted Luciferin Detection Reagent (100 μL) to each well of a 96-well plate. Mixed briefly on a plate shaker, incubated samples for 15 minutes and measured luminescence. Raw luminescent numbers were quantified to total glutathione (μM) by using a total glutathione standard curve.

### DILI Biomarkers

<u>Keratin 18 (K18):</u> levels were quantified in human model effluent samples from both the hepatocyte and vascular channels using the M65 EpiDeath CK18 ELISA Kit (DiaPharma, Detroit, MI, USA). The assay was run following the vendor protocol using a standard curve ranging from 0-5,000 U/L. Additionally, a standard curve ranging from 0-500 U/L was generated independently for quality control.

<u>Alpha Glutathione S-Transferase (α-GST)</u>: levels were quantified in human model effluent samples from the upper channel using an ELISA kit (DiaPharma). The assay was run following the vendor protocol using a standard curve ranging from 0-64 µg/L. Additionally, a standard curve ranging from 0-50 µg/L was generated independently for quality control.

<u>Mir-122:</u> Isolation of micro-RNAs was performed on upper and lower chamber eluates using Qiagen’s (Hilden, Germany) miRNA isolation kit for serum and plasma. A total volume of 50 μL was used for isolation. Isolation was performed according to manufacturer’s protocol using a 56.5 pM oligonucleotide spike-in control. Samples were eluted with 14 μL RNase-free water. Reverse transcription was performed using Thermo Fisher Scientific’s TaqMan^TM^ MicroRNA Reverse Transcription Kit according to the manufacturer’s protocol. A total reaction volume of 15 μL was used. Real-time PCR was performed using Thermo Fisher Scientific’s TaqMan® Micro-RNA Assays according to manufacturer’s protocol. A total reaction volume of 10 μL was used. Normalized ΔΔCt was used for fold-change analysis.

<u>Cytokines</u>: Release of IL-6, MCP-1, and IP-10 from the vascular channel was quantified using U-PLEX biomarker human assays (Meso Scale Diagnostics, Rockville, MD, USA) and the assays were performed according to the protocol provided by the vendor.

### Gene expression analysis

Gene expression levels were analyzed using 2-step PCR. Total RNA isolation was performed using the RNeasy 96 Kit (Qiagen) or PureLink® RNA Mini Kit (Invitrogen) following the vendor protocol. To prepare cDNA, a combination of SuperScript VILO MasterMix or SuperScript^TM^ IV (Invitrogen) and water was added to the isolated RNA samples and run on iCycler (BioRad) using the SuperScript VILO protocol or SimpliAmp Thermocycler (Applied Biosystems, Waltham, MA, USA) using the SuperScript™ IV First-Strand Synthesis System protocol. Subsequent addition of a mixture of TaqMan Universal PCR MasterMix or TaqMan Fast Advanced MasterMix (Thermo Fisher), water, and the appropriate probe was added to the cDNA samples and run on ViiA7 Real-Time PCR System (Thermo Fisher) or QuantStudio™ 3 Real-Time PCR System (Applied Biosystems) for qPCR analysis. Relative gene expression levels were calculated using ddCt method.

### CYP450 enzyme activity measurement

CYP450 enzyme activity was determined using prototypical probe substrate compounds: phenacetin, midazolam, and bupropion (Sigma) as a cocktail or a single substrate, and cyclophosphamide and testosterone (Sigma) as a single substrate. For the cocktail substrate, a mixture of phenacetin at 30 µM, midazolam at 3 µM, and bupropion at 40 µM final concentration respectively, was prepared in serum-free medium. For the single substrate, phenacetin at 100 µM, cyclophosphamide at 1 mM, and testosterone at 200 µM final concentration respectively, was prepared in serum-free medium. Enzyme activities were measured at Day 0, 3, 7, and 14 of hepatocyte cultures after a 30-min for testosterone, 1-hour and 2-hours incubation for the rest of the probe substrates under a flow rate of 200~250 µL/hr of flow rate. The control condition was tested with hepatocytes in static sandwich monoculture plates using the same concentration of substrate and duration of incubation, using 500 µL in 24-well plate format. The reaction was stopped using acetonitrile with 0.1% formic acid, and formation of metabolites were measured using LC-MS at AstraZeneca or at In Vitro ADMET Laboratories, Inc. (IVAL).

### APAP metabolite quantification

After APAP exposure to human Liver-Chips, effluent metabolite levels were quantified using LC-MS analysis. Relative quantitation was performed by first generating a standard curve using six concentrations of APAP standards: 0.78, 1.56, 3.12, 6.24, 12.48 and 24.96 mg/mL. These concentrations were plotted versus LC-MS peak area to generate a standard curve and subsequent linear regression equation: Y=0.5046.3*X. Effluent samples from Day 20 of APAP treatment were then analyzed, and the concentration of APAP present was interpolated using the standard curve regression: X=Y/5046.3*D, where X is the concentration of APAP, Y is the peak area from LC-MS, and D is the dilution factor. APAP-Glucuronide (APAP-Glu), the primary metabolite of APAP, was not quantified using a standard curve. Rather, the LC-MS peak area of APAP-Glu was reported corresponding to each sample.

### Testing of JNJ-2 in rats

Following the discovery of liver fibrosis in a 14-day male rat study with JNJ-2, a mechanistic study was performed with the same compound at the high dose, three time points (3, 7, and 14 days), 12 male rats per group (6 euthanized at the end of treatment, and 6 after a 14-day recovery period). Regarding histology procedures, the liver was sampled. Briefly, 5 serial sections were prepared. One was stained with hematoxylin and eosin (H&E) kit. The 4 others were submitted to histochemical and immunohistochemical stains. Collagen was detected with the Van Gieson kit (Merck, Darmstadt, Germany), and reticulin, with the Reticulum kit (Sigma-Aldrich). A monoclonal mouse anti-human αSMA antibody (Dako, Denmark) was incubated for 2 hours at room temperature after antigen retrieval, endogenous peroxidase block and with normal goat serum block, and revealed with the detection kit Vectastain ABC Elite (Vector Labs, Burlingame, CA, USA) and the chromogen DAB (Dako). A monoclonal mouse anti-rat CD68 (ED1) antibody (Serotec, Raleigh, NC, USA) followed the same protocol.

### Testing of JNJ-3 in dogs

JNJ-3 (in 20% hydroxypropyl-β-cyclodextrin) was administered once daily for 14 days to 3 male beagle dogs per group at doses of 15 and 65 mg/kg. Mortality, clinical observations, body weight, food consumption, clinical pathology, gross necropsy and microscopic examination of selected tissues, and toxicokinetics were evaluated. Dogs were fasted overnight prior to blood collection for measuring clinical pathology parameters (including AST and ALT) after 6 and 14 days of dosing. Histology slides (hematoxylin and eosin staining and Mason’s Trichrome staining) of the liver tissues were prepared and evaluated microscopically. The quantification of JNJ-3 was conducted using a qualified liquid chromatographic-triple quadrupole mass spectrometric (LC-MS/MS) procedure.

### Semi-quantitation of TAK-875 metabolites in eluates from human Liver-Chip

#### Incubation of ^14^C-TAK-875 with human liver microsomes in the presence of NRS and UDPGA/UDPAG

The goal of the *in vitro* incubation is to generate ^14^C-TAK-875 acyl glucuronide and ^14^C-M1 metabolite for quantitation of these 2 metabolites in incubates from human Liver-Chip model. Human liver microsomes (HLM, 1 mg/mL, BD Gentest, Franklin Lakes, NJ, USA) was pre-incubated with 10 µM ^14^C-TAK-875 at 37℃ for 3 minutes. The NADPH regenerating system and uridine 5’-diphosphoglucuronic acid (UDPGA, 5 mM) : uridine 5’-diphospho-N-acetylglucosamine (UDPAG, 1mM) were added to initiate the reaction at the end of pre-incubation. The reaction mixtures were further incubated for 60 minutes. Incubates in the absence of radiolabeled TAK-875 served as calibration matrices for samples from Liver-Chip. The reaction was quenched with 5 volumes of acetonitrile:isopropyl alcohol/1:1 fortified with 0.1% formic acid and ammonium formate (500 mM, pH 3.0) to stabilize the acyl glucuronides. The mixture was vortexed mixed and sonicated prior to centrifugation at 3000 g for 10 minutes at 4°C. The resulting supernatants were dried under nitrogen. The dried residues were suspended in 300 μL of acetonitrile:water:isopropyl alcohol/1:2:1 fortified with 0.1% formic acid. The suspension was filtered through 0.45 μm Nylon filters for LC/RAD/MS analysis.

#### Preparation of Samples for quantitation by calibration of MS response

An equal volume of ^14^C-TAK-875 HLM incubate and sample matrices (vehicle-treated) from Liver-Chip were mixed prior to analysis. Similarly, samples from Liver-Chip following treatment with cold TAK-875 at 10 μM for ~ 2weeks were mixed with the HLM matrices (without ^14^C-JNJ-TAK-875) prior to analysis.

The unchanged drug, M1 and acyl glucuronide metabolites of TAK-875 in eluates from Liver-Chip were quantified using the peak areas from ^14^C and MS of HLM incubate according to the equation as follows:

(^14^C-Peak Area/Peak area of same component from MS ionization of ^14^C sample)* Peak area of same component from MS ionization of Emulate samples

### Image analysis

<u>CLF:</u> Both a brightfield and a fluorescence image was taken from each field of view. Both images were de-noised using median filtering, locally enhanced in contrast using the CLAHE algorithm, and thresholded to extract bright regions which corresponded to the biliary canaliculi (channel 1) and to CLF-labeled regions (channel 2), respectively, using Matlab (MathWorks, Natick, MA, USA). CLF-labeled regions that most likely to corresponded to bile canaliculi were automatically identified based on geometric criteria and retained only if they co-localized with a bile canaliculi signal in channel 1. Specifically, the main criteria used to detect canalicular geometry were: (1a) Circularity is lower than 0.5 and eccentricity is greater than 0.8 (jagged, elongated canaliculi), or (1b) Eccentricity is greater than 0.8 and solidity is greater than 0.7 (smooth, elongated canaliculi), and (2) Total size is smaller than 70 μm^2^ and greater than 7 μm^2^ (excluding noise and stained cell debris). These criteria were determined empirically using a small test set of images and then applied to all images for the analysis. Circularity, eccentricity, and solidity were computed using Matlab built-in functions. The detected areas hence correspond to CLF-containing bile canaliculi. The data for treated and control samples were quantified by computing the area percentage of CLF-labeled bile canaliculi within each field of view (FOV). Because of the non-normal distribution of these data, statistical analysis was performed using the non-parametric Wilcoxon rank sum test. The total number of analyzed FOVs from 2 chips each were N=8 FOVs for 30 μM bosentan and N=17 FOVs for vehicle-treated samples (0.1% DMSO).

<u>BSEP area quantification:</u> The fluorescent channel images were first de-blurred with Olympus cellSens software. Using ImageJ, they were then histogram adjusted to remove background, adaptive thresholded to extract fluorescent regions (a plugin developed by Qingzong Tseng using the adaptive threshold method of the OpenCV library), and analyzed using the built-in 3D Object Counter function with an area cutoff value to remove noise. N=3 FOVs from each chip were analyzed for vehicle (0.1% DMSO) and 30 µM bosentan conditions.

<u>Lipid accumulation quantification:</u> The stained chips were imaged using the TRITC filter, and were de-blurred with Olympus cellSens software. Using ImageJ-Fiji, the fluorescent channel images were histogram adjusted to remove background, followed by fluorescence quantification using the Integrated Density calculation function. N=5 FOVs per chip were analyzed for all conditions: vehicle (0.1% DMSO), FIAU dosing groups, and MTX dosing groups.

<u>Kupffer cell and activated stellate cell count:</u> The number of CD68- or α-SMA-positive cells in the vascular channel in each group was counted in each FOV (360 mm^2^) and quantified.

<u>α-SMA intensity quantification</u>: the fluorescent channel (588) was first background subtracted and processed using the median filter of Image J. After processing, the signal was thresholded using adaptive thresholding function of Image J (Otsu method) to extract fluorescent regions and to measure the relative intensity of the fluorescent signal. N=4 to 6 FOVs per chip from each chip were analyzed. All results were normalized to the relative control condition (vehicle, DMSO 0.1%) and reported as fold increment in respect to control.

### Statistical analysis

As indicated in the figure legends, one-way ANOVA, Sidak’s and Dunnett’s multiple comparisons tests were used for parametric data and the Mann-Whitney U test or Kruskal-Wallis tests was used for nonparametric data. All statistical analyses were performed using Prism 7 (GraphPad).

## Supplementary Figures

**Fig. S1.**
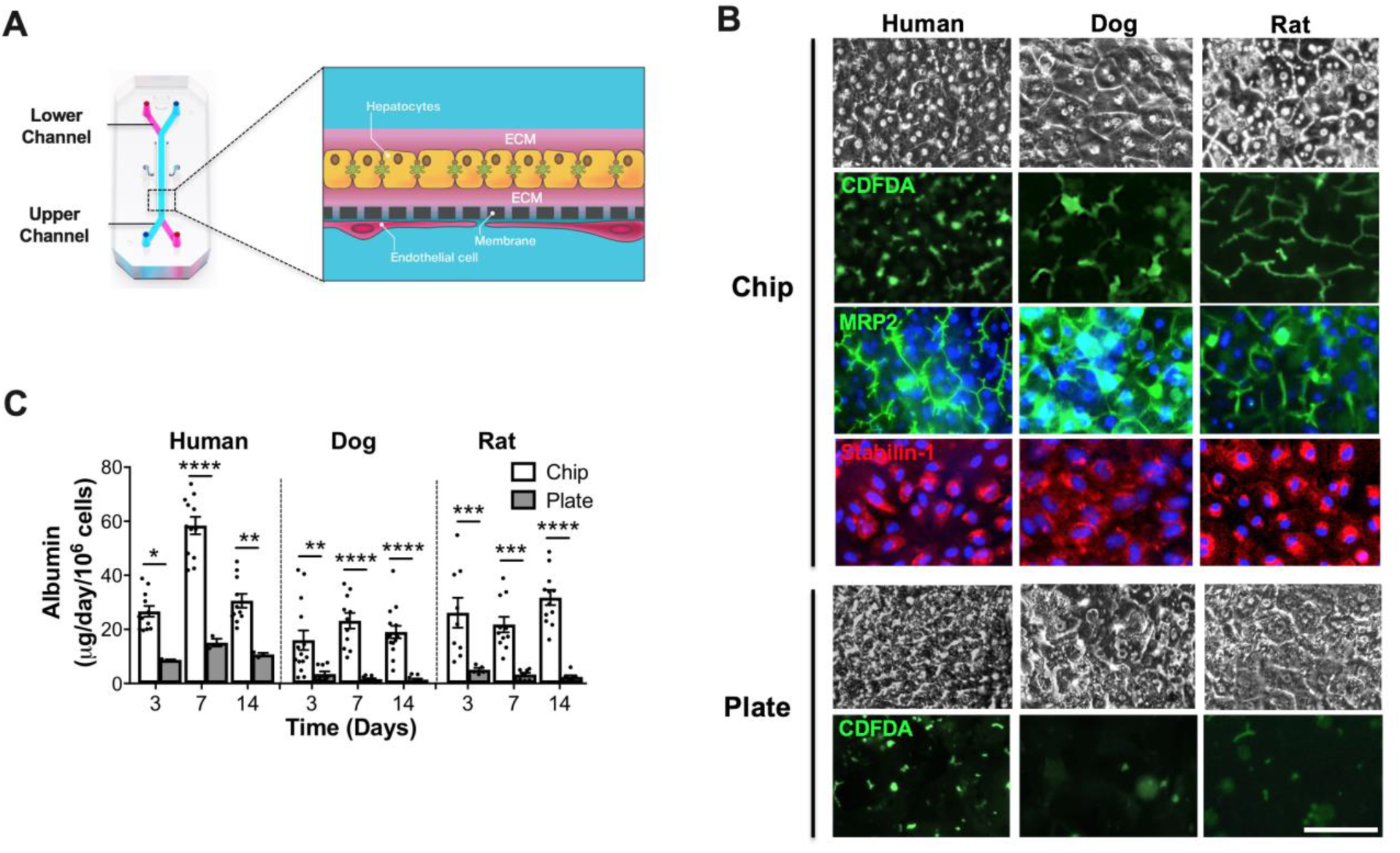
Morphology and functionality of species-specific dual-cell Liver-Chips. (**A**) Schematic of the dual-cell Liver-Chip that recapitulates complex liver microarchitecture. Primary hepatocytes in the upper channel in ECM sandwich format and LSECs on the opposite side of the same membrane in the lower vascular channel. (**B**) Representative images of hepatocytes (bright-field), CDFDA (green) to visualize bile canaliculi in hepatocytes, MRP2 (green and DAPI in blue) in hepatocytes, and stabilin-1 (red and DAPI in blue) in LSECs after 14 days of culture in human, dog, and rat Liver-Chips and sandwich monoculture plates. Scale bar, 100 µm. (**C**) Albumin secretion in human, dog, and rat Liver-Chips over 2 weeks compared to static sandwich monoculture plates. Sidak’s multiple comparisons test (n=3~14 independent Chips, n=3~9 independent wells in plate). *P < 0.05, **P < 0.01, ***P < 0.001, ****P < 0.0001. Error bars present mean±SEM.

**Fig. S2.**
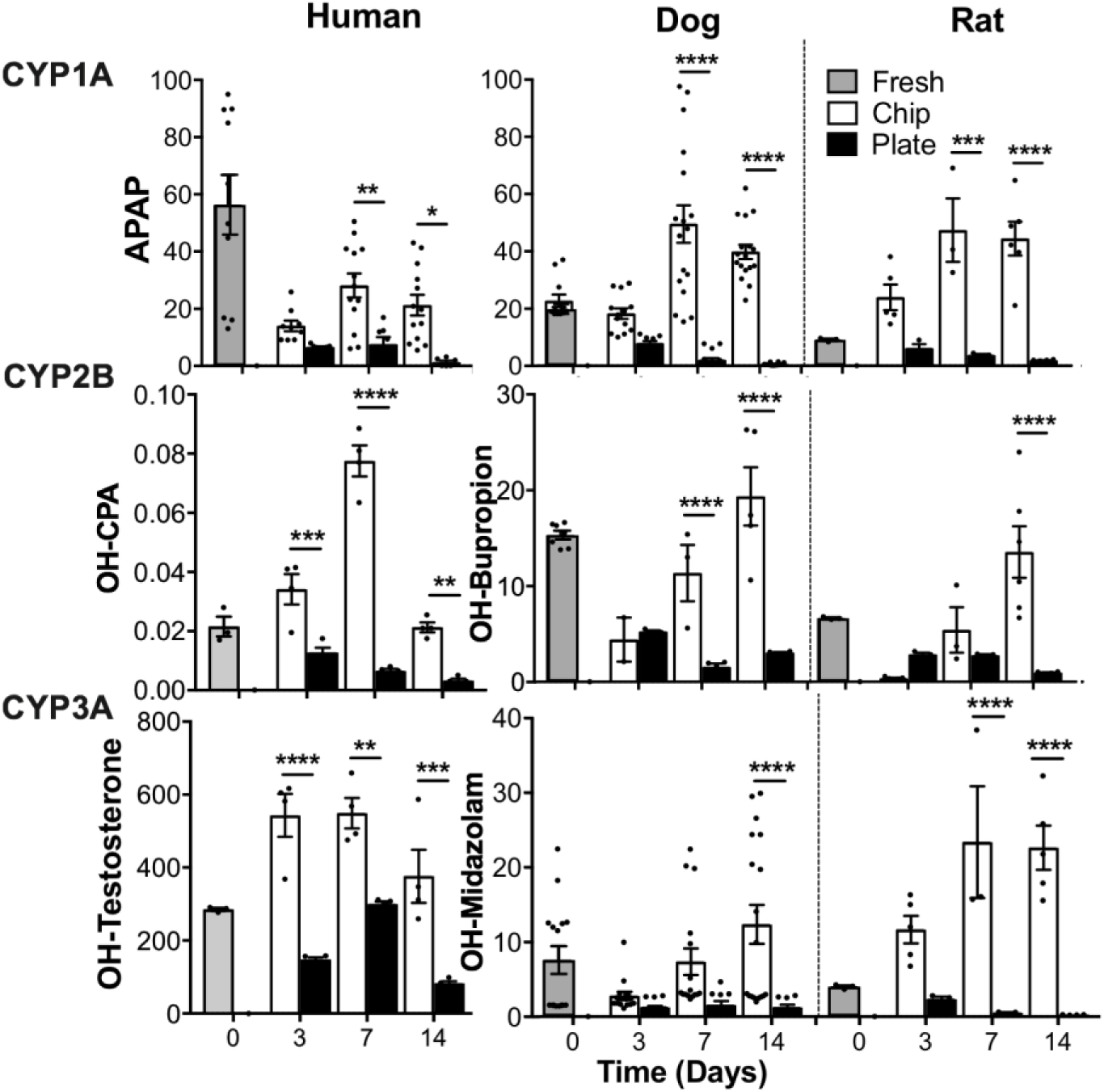
Cytochrome P450 enzyme activity in species-specific dual-cell Liver-Chips. Cytochrome P450 enzyme activity in human, dog, and rat Liver-Chips compared to conventional sandwich monoculture plates and fresh hepatocyte suspension over 2 weeks using a cocktail (for dog and rat) or single (for human) probe substrate. Unit: pmol/min/10^6^ cells. Sidak’s multiple comparisons test (n=3~17 independent chips). *P < 0.05, **P < 0.01, ***P < 0.001, ****P < 0.0001. Error bars present mean±SEM.

**Fig. S3.**
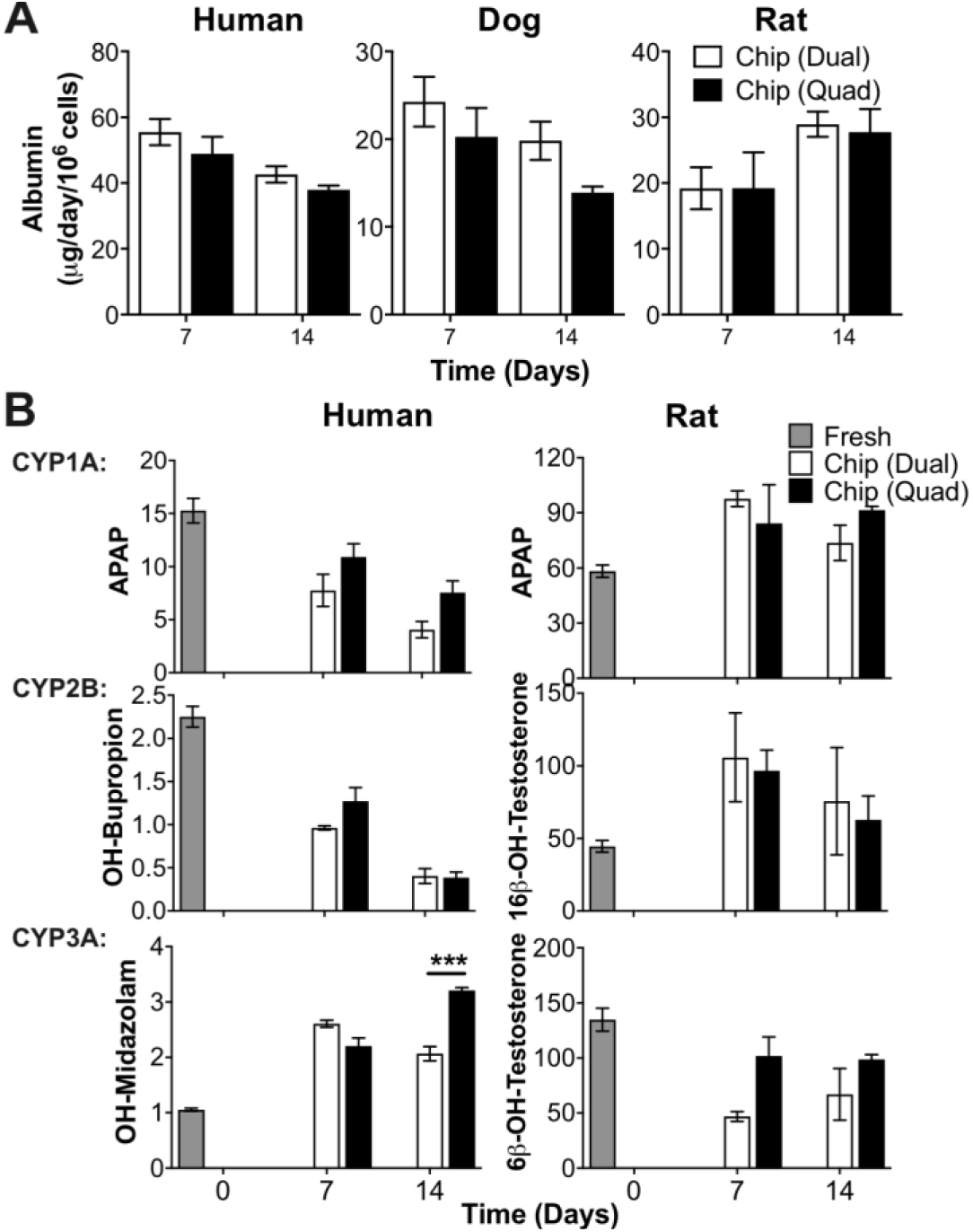
Comparison of hepatic functionalities between dual- and quadruple-cell Liver-Chips. (**A**) Comparison of albumin secretions between dual- and quadruple-cell Liver-Chips from three species models. (**B**) Comparison of CYP450 enzyme activities between dual- and quadruple-cell Liver-Chips from human and rat models. Sidak’s multiple comparisons test (n=3~4 independent chips). ***P < 0.001. Error bars present mean±SEM.

**Fig. S4.**
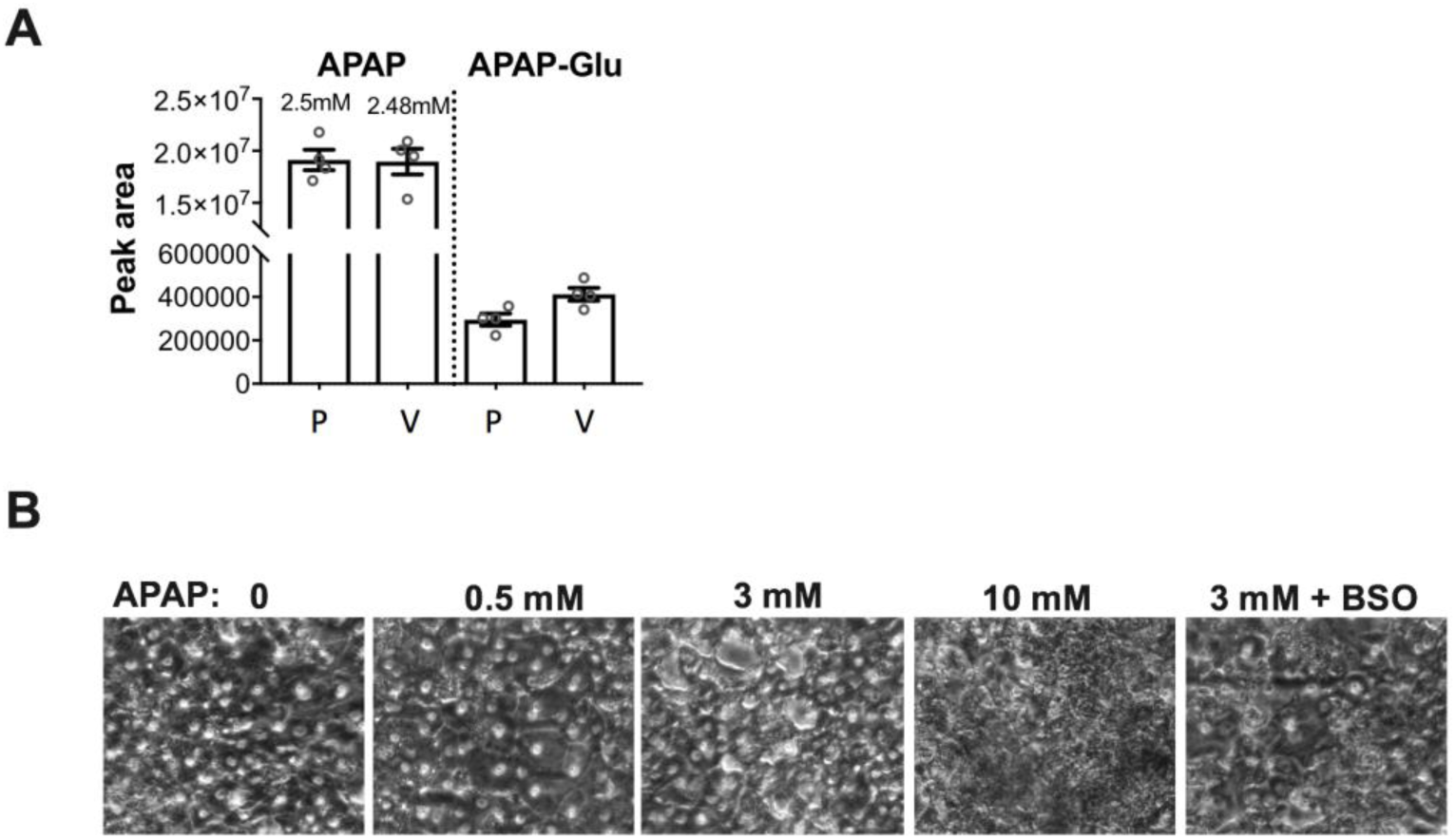
Detection of glucuronide metabolites of APAP and hepatocellular injury using quadruple-cell human Liver-Chips. (**A**) APAP glucuronide metabolites formation from upper parenchymal (P) and lower vascular (V) channels after APAP treatment at 3 mM for 20 days from human Liver-Chips. (n=4 independent chips). (**B**) Representative bright-field images of hepatocytes after daily administration of APAP at 0.5, 3, and 10 mM and co-administration of APAP 3 mM and 200 μM of buthionine sulfoximine (BSO) for 7 days in human Liver-Chips

